# Insights into Human Epileptogenesis with Proteomic Profiling

**DOI:** 10.1101/2024.01.01.573812

**Authors:** Najing Zhou, Yang Di, Di Zhang, Wei Gang, Huiran Zhang, Yi Yuan, Aitao Zhang, Jie Dai, Xiaona Du, Wenling Li, Hailin Zhang

## Abstract

Epilepsy affects millions globally, and drug-resistant epilepsy remains a challenge. Molecular mechanisms underlying epilepsy remain elusive. Protein profiling through proteomics offers insight into biomarkers and therapeutic targets. Human brain tissue from epilepsy surgeries was analyzed using data-independent acquisition (DIA) proteomics. Samples were categorized into Core (epileptogenic focus), Border (marginal excision tissue), and Nonepileptic control groups. Differential expression proteins (DEPs) were identified and shared proteins were analyzed. 163 DEPs were identified which may has potential roles in the initiation of epileptic electrical firing, 412 DEPs which indicating the difference between epilepsy and Nonepilepsy patients and 10 DEPs consistently altered in Core which indicating potential roles in epileptogenesis. Notably, P35754/GLRX, O75335/PPFIA4, and Q96KP4/CNDP2 were consistently expressed differently in all group pairs. From validation experiments, the expression of Kv3.2 significant reduced in the Core group compare to border group by immunohistochemistry and knockdown of Kv3.2 increased seizure susceptibility and altered neuronal excitability through our cellular and animal experimentation.

## Introduction

Epilepsy is a chronic neurological disorder characterized by recurrent seizures, affecting approximately 50 million people worldwide(World Health Organization, 2023). Despite the availability of over 30 FDA-approved anti-epileptic drugs, approximately 30% of cases will develop drug-resistant or refractory epilepsy(West et al, 2015). Among these cases, temporal lobe epilepsy (TLE) is the most common, with about 40% of TLE cases becoming drug-resistant(Fisher et al, 2014). Epilepsy surgery offers the most promising chance for a cure in patients with medically refractory epilepsy, with approximately 60% of TLE patients achieving freedom from seizures when surgical intervention is available(West et al, 2015). Although epilepsy surgery is generally considered a safe and effective treatment option for patients with medically refractory epilepsy. However, as with any surgical procedure, there are potential risks and complications associated with epilepsy surgery, including infection, bleeding, stroke, and cognitive or neurological deficits(Tellez-Zenteno et al, 2005; Wiebe et al, 2001).

Despite extensive research, the underlying molecular mechanisms of epilepsy remain poorly understood(Kirchner et al, 2020). Understanding molecular mechanism of epileptogenesis is a challenging but a crucial step in finding new drug target for treatment of epilepsy. Proteins, as the major functional units of cells, play a critical role in the pathogenesis of epilepsy. Protein profiling using proteomics methods has emerged as a promising approach to identify potential biomarkers and therapeutic targets for epilepsy(Bruxel et al, 2021; Grant & Blackstock, 2001).

In recent years, there has been a growing interest in using human samples obtained from epilepsy surgery to investigate the protein alterations associated with epileptogenesis (Barker-Haliski et al, 2015; Gorter et al, 2015). Surgical specimens provide an opportunity to directly analyze the affected brain tissue and identify changes in protein expression, post-translational modifications, and protein-protein interactions that are specific to epilepsy.

Proteinomics methods such as mass spectrometry-based proteomics, protein microarrays, and antibody-based assays can be applied to comprehensively characterize the proteome of epileptic brain tissue(Cravatt et al, 2007). These methods have the potential to identify novel biomarkers of epilepsy, provide insights into the molecular pathways involved in the development of epilepsy, and facilitate the discovery of new therapeutic targets. Recent studies using proteinomics methods have identified alterations in the expression of various proteins in the brain tissue of patients with epilepsy, including changes in the expression of ion channels, synaptic proteins, and enzymes involved in energy metabolism(Banote et al, 2021; Chen et al, 2021; Do et al, 2020; Gorter et al, 2006). These studies suggest that alterations in the expression of specific proteins may contribute to the development and progression of epilepsy. However, the results of many studies investigating protein alterations in epilepsy vary greatly due to differences in experimental designs, sample sources of different patients and sites, and various proteome detection techniques. This variability can make it challenging to identify key molecular targets for the development and treatment of epilepsy. Standardized protocols and rigorous quality control measures are needed to improve the reproducibility and reliability of proteomics data in epilepsy research.

This study aims to utilize DIA proteinomics method(Pappireddi et al, 2019) to analyze human brain tissue samples obtained from epilepsy surgery and identify protein alterations associated with epileptogenesis. The results of this study may lead to the development of new diagnostic and therapeutic strategies for epilepsy.

## Results

We conducted a multigroup proteomic analysis of temporal lobectomy samples obtained from 9 patients with refractory medial temporal lobe epilepsy and 8 patients with non-epileptic and non-tumorous conditions using DIA method. The epileptic samples were further divided into two groups: Core group (the region of epileptogenic focus) and Border group (the marginal area of excision tissue). The non-epileptic samples were used as Nonepileptic control group. The clinical data of the patients are presented in Table 1-2. Bioinformatics and statistical analysis were performed among these three groups. Our interpretation of the data from these three groups is that Core and Border samples from patients who underwent epileptic surgery are associated with epilepsy, and the differences between them may indicate potential mechanisms for the initiation of epileptic electrical firing. On the other hand, the differences between Core and Nonepileptic, between Border and Nonepileptic groups likely represent general mechanisms of epileptogenesis. Based on this thinking, we conducted a focused study on the differentially expressed proteins (DEPs) that showed consistent changes in expression levels (either increase or decrease) between the Core *vs.* Border and between Core/Border *vs.* Nonepileptic. This allowed us to identify a subset of proteins that were consistently dysregulated within the epileptic zone, as well as in the whole epileptic tissue, regardless of the location within the tissue.

**Table 1:** The clinical data of epilepsy patients.

|  | Gender | Age | Time from seizure onset | Diagnosis | Pathologic diagnosis | Surgery details |
| --- | --- | --- | --- | --- | --- | --- |
| 1 | Female | 14 | 4 years | 1. Symptomatic epilepsy<br>2. Degeneration of left temporal neurons | FCDIIa<sup>A</sup> | Left ATL<sup>B</sup> |
| 2 | Female | 37 | 5 years | Symptomatic epilepsy | FCDIIIb | Right ATL |
| 3 | Male | 21 | 6 years | 1. Epilepsy<br>2. Central nervous system infection<br>3. Left hippocampal sclerosis | FCDIIIa | Left ATL |
| 4 | Male | 38 | 2 months | 1. Symptomatic epilepsy<br>2. Left medial temporal structure FCD | FCDIIIa | Left ATL |
| 5 | Female | 24 | 10 years | Epilepsy | FCDIIIb | Right ATL + right temporal base eclampsia resection |
| 6 | Male | 43 | 2 years | 1. Left anterior temporal focal cortical dysplasia<br>2. Symptomatic epilepsy | FCD IIIb | Left ATL |
| 7 | Male | 29 | 6 months | Symptomatic epilepsy | Meningeal angiomatosis | Right ATL |
| 8 | Female | 27 | 24 years | 1. Symptomatic epilepsy<br>2. Right hippocampal sclerosis | FCDIIIa | Right ATL |
| 9 | Male | 8 | 2 years | Epilepsy | FCDIIIa | Left ATL |

Footnote: <sup>A</sup>FCD, Focal cortical dysplasia;
<sup>B</sup>ATL, Anterior temporal lobectomy.

**Table 2:** The clinical data of non-epilepsy patients.

|  | Gender | Age | Basic information | Diagnosis | Surgery details |
| --- | --- | --- | --- | --- | --- |
| 1 | Male | 59 | Unconscious after being hit by a car. Diabetic | Left temporal contusion, left temporal parietal subdural hematoma, traumatic subarachnoid hemorrhage | Removal of left temporal laceration hematoma |
| 2 | Male | 36 | Unconscious after a fall on an electric bike | Cerebral contusion, traumatic subarachnoid hemorrhage, right frontotemporal parietal subdural hematoma, left parietal epidural hematoma | Right temporal brain contusion removal + right frontotemporal parietal subdural hematoma + right frontotemporal parietal craniotomy decompression |
| 3 | Male | 64 | Poor speech, confusion, high blood pressure for 20 years | Cerebral hemorrhage in left basal ganglia | Removal of hematoma from left basal ganglia hemorrhage + Decompression of bone flap |
| 4 | Male | 35 | Sudden collapse with unconsciousness | Cerebral hemorrhage in left basal ganglia | Removal of hematoma from left basal ganglia hemorrhage + Decompression of bone flap |
| 5 | Female | 68 | High blood pressure for 10 years | Massive hemorrhage in right basal ganglia, frontotemporal lobe, cerebral hernia, subarachnoid hemorrhage | Removal of hematoma from left basal ganglia hemorrhage + Decompression of bone flap |
| 6 | Female | 57 | Sudden headache with unconsciousness, high blood pressure for 20 years | Cerebral hemorrhage in left basal ganglia | Removal of hematoma from left basal ganglia hemorrhage + Decompression of bone flap |
| 7 | Male | 51 | Sudden hemiplegia, high blood pressure for 6 years | Cerebral hemorrhage in left basal ganglia | Removal of hematoma from left basal ganglia hemorrhage + Decompression of bone flap |
| 8 | Male | 37 | Sudden right limb weakness with unconscious nausea and vomiting after drinking alcohol | Cerebral hemorrhage in left basal ganglia | Removal of hematoma from left basal ganglia hemorrhage + Decompression of bone flap |

*Missing value imputation and qualitative analysis of identified proteins* To ensure the validity and accuracy of subsequent bioinformatics and statistical analyses, we only included proteins with quantitative values detected in more than 50% of samples from each group. Although we used DIA, a proteomic assay that significantly improved the consistency of protein detection, missing values (or not available data points) can still negatively affect subsequent proteomic data analyses. Therefore, we adopted the method proposed by Wang et al., and imputed missing values using the local similarity method Seq-KNN(Wang et al, 2020).

We identified a total of 6476 proteins and peptides in Nonepileptic control group, 6401 in Border group, and 6427 in Core group. Among these proteins, 6220 were common to all three groups. In addition, 89 proteins were shared between Core group and Border group, 95 proteins were shared between Core group and Nonepileptic control group, and 74 proteins were shared between Border group and Nonepileptic control group. Furthermore, 86 proteins were unique to Nonepileptic control group, 17 to Border group, and 22 to Core group. The Venn diagram is presented in Figure 1A.

**Figuure 1.**
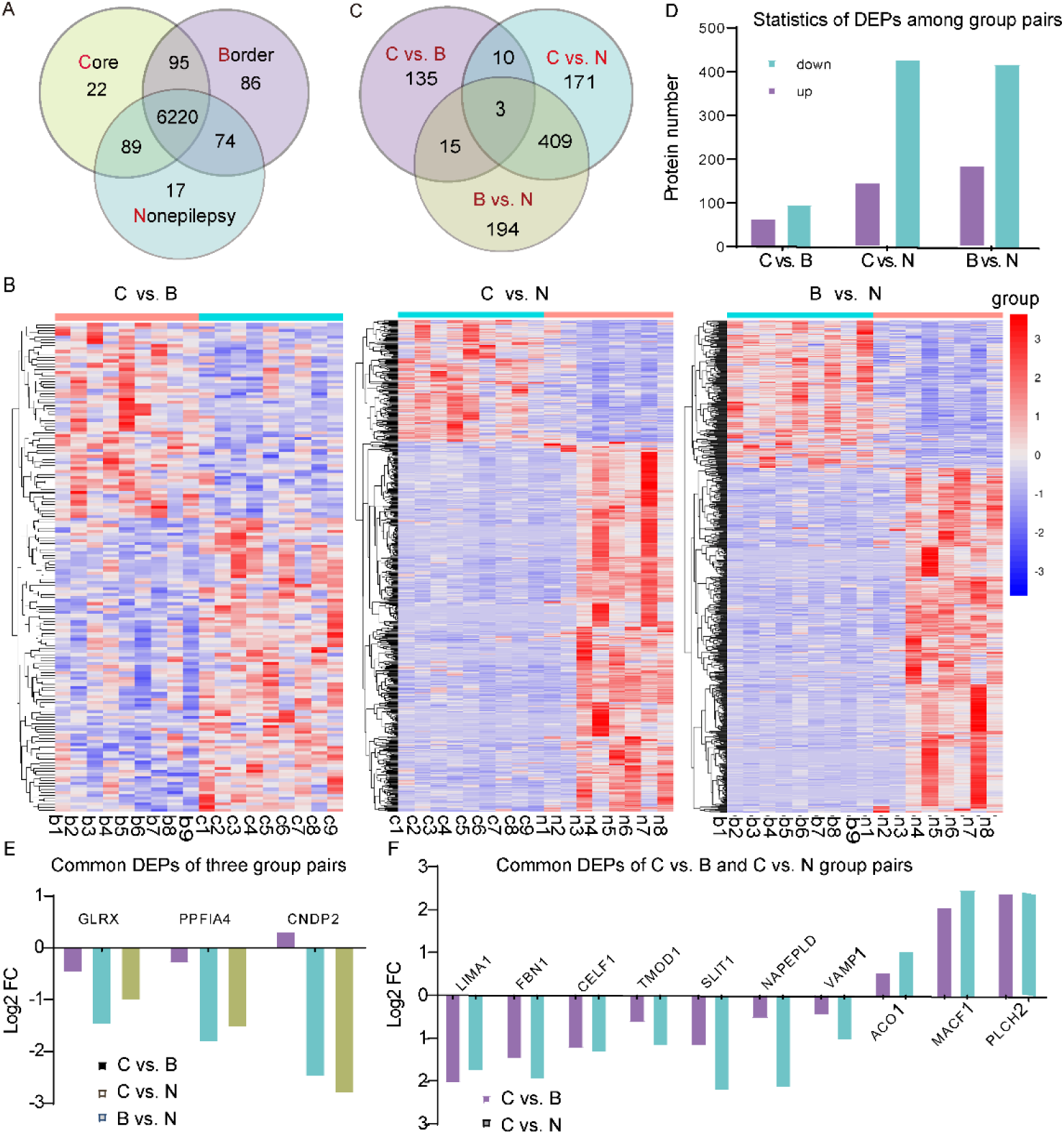
Statistics of protein expression and different expressed proteins between any two groups of Core, Border and Nonepilepsy with human samples. A, Venn statistics of protein expression of all three groups and their common proteins. B, Cluster analysis of different expressed proteins (DEPs) between any two groups of Core, Border and Nonepilepsy. C, Venn statistics of DEPs between any two groups of Core, Border and Nonepilepsy. D, Statistics of DEPs expression changes of varied group pairs. E, 3 common DEPs of three group pairs. F, 10 common DEPs of core vs. Border and Core vs. Nonepilepsy group pairs. Notes: C, Core; B, Border; N, Nonepilepsy.

Cluster analysis shown in Figure 1 B revealed a distinction between Core and Border group, and between Core or Border group and Nonepileptic control group.

*Differential expression proteins and peptides (DEPs) between groups* The *t*-test was used to compare Border group with Nonepileptic control group or Core group with Nonepileptic control group, and a paired *t*-test was used for Core group with Border group. The criteria for identifying DEPs were based on the fold change of average value being more than 1.2 or less than 0.83 for Core *vs.* Border group pair, more than 2 or less than 0.5 for Border *vs.* Nonepileptic control group pair and Core *vs.* Nonepileptic control group pair, as well as a *p*-value less than 0.05.

We identified 163 DEPs in Core group compared with Border group, of which 65 were up-regulated and 98 were down-regulated. Between Core and Nonepileptic control groups, there were 593 DEPs, of which 146 proteins were up-regulated, and 447 proteins were down-regulated. In addition, we found 621 DEPs between Border group and Nonepileptic control group, of which 186 proteins were up-regulated, and 435 proteins were down-regulated. (Figure 1C and D). The DEPs were listed in supplement Datasets S1.

Upon further analysis, we identified three DEPs that were consistently differentially expressed in all three group pairs (see Figure 1E). These three proteins are P35754/GLRX, O75335/PPFIA4, and Q96KP4/CNDP2. Notably, all three proteins showed significant abundance levels.

In the case of P35754/GLRX and O75335/PPFIA4, their expression was found to be lower in Border group compared to Nonepileptic control group, and the lowest expression was observed in Core group (Figure 1E). This finding suggests a potential role for these proteins in epileptogenesis. P35754/GLRX functions as a glutathione-disulfide oxidoreductase in the presence of NADPH and glutathione reductase(Knoops et al, 1999). On the other hand, O75335/PPFIA4 is implicated in regulating the disassembly of focal adhesions and localizing receptor-like tyrosine phosphatases type 2A at specific sites on the plasma membrane (Xie et al, 2020).

For Q96KP4/CNDP2, its expression was lowest in Border group and slightly higher in Core group compared to Border group. Q96KP4/CNDP2 has the ability to hydrolyze various dipeptides and activate the mitogen-activated protein kinase (MAPK) pathway (Zhang et al, 2014). An increased level of CNDP2 activates the p38 and JNK MAPK pathways, leading to cell apoptosis, while a lower level of CNDP2 activates the ERK MAPK pathway, promoting cell proliferation (Zhang et al, 2014). These findings suggest that P35754/GLRX, O75335/PPFIA4, and Q96KP4/CNDP2 may play important roles in the pathogenesis of epilepsy.

Next, our analysis focused on identifying common proteins with consistent changes in expression levels between the Core *vs.* Border group pair and the Core *vs.* Nonepileptic control group pair. We identified a total of 10 DEPs that showed consistent alterations. Among these 10 DEPs, 7 were expressed at lower levels in Core group compared to both Border and Nonepileptic control groups, while 3 were expressed at higher levels in Core group compared to both Border and Nonepileptic control groups (Figure 1F).

One of these DEPs is NAPEPLD, which is known to be involved in controlling N-acyl-phosphatidylethanolamine (NAPE) homeostasis in dopaminergic neuron membranes and regulating neuron survival (Palese et al, 2019). PLCH2 is a phosphatidylinositol-specific phospholipase C enzyme that is very sensitive to calcium, and can cleave PtdIns(4,5)P2 to generate second messengers inositol 1,4,5-trisphosphate and diacylglycerol. PLCH2 may be important for formation and maintenance of the neuronal network in the postnatal brain (Kanemaru et al, 2010). The presence of these 10 DEPs with consistent expression changes suggests that they may play essential roles in the mechanisms underlying epileptogenesis.

Finally, we conducted an analysis to identify common proteins with consistent changes in expression levels between the Core/Border *vs.* Nonepileptic control pairs. We found a total of 409 DEPs. When combined with the three differential proteins shared by all three groups, there were a total of 412 proteins that showed significant differences between the epileptic Core/Border groups and the Nonepileptic control samples (Datasets S2).

These 412 DEPs represent the most prominent differences between individuals with epilepsy and those without epilepsy. They may serve as intrinsic pathogenic factors involved in epileptogenesis.

*Gene Ontology (GO) and KEGG pathway analysis of DEPs* We performed Gene Ontology (GO) and KEGG pathway analyses on the previously mentioned DEPs for each comparison, including the Core *vs.* Border, Core *vs.* Nonepileptic, and Border *vs.* Nonepileptic pairs. The enriched cellular component (CC), biological process (BP), and molecular function (MF) components, as well as the enriched KEGG pathways, were summarized in supplement Figure S1. Enriched KEGG pathways are associated with these DEPs presented in Datasets S3 and the most significant pathways are presented in Figure 2. The network of “Regulation of actin cytoskeleton” and “Endocytosis” signal pathways as typical examples demonstrating the differences of pathway-related signal networks between group pairs are shown in Figure 3.

**Figure 2:**
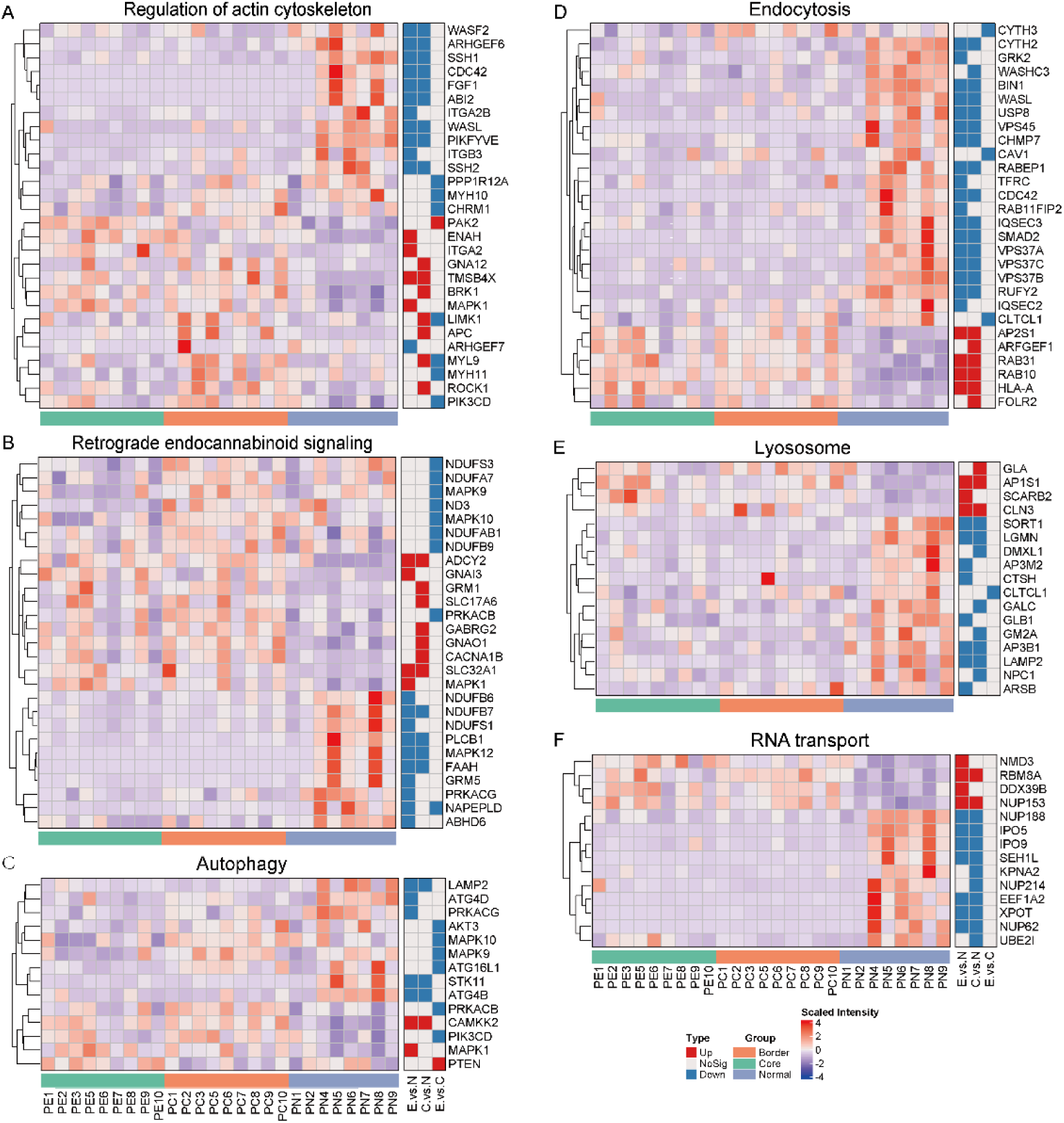
Cluster analysis of the DEPs of a few top different KEGG pathways from three group pairs. A. DEPs of regulation of actin cytoskeleton pathway. B. DEPs of retrograde endocannabinoid signaling pathway. C. DEPs of autophagy pathway. D. DEPs of dopaminergic synapse signaling pathway. E. DEPs of apoptosis pathway. F. DEPs of RNA transport pathway. G. DEPs of endocytosis pathway. H. DEPs of lysosome pathway. I. DEPs of adherent junction pathway.

**Figure 3.**
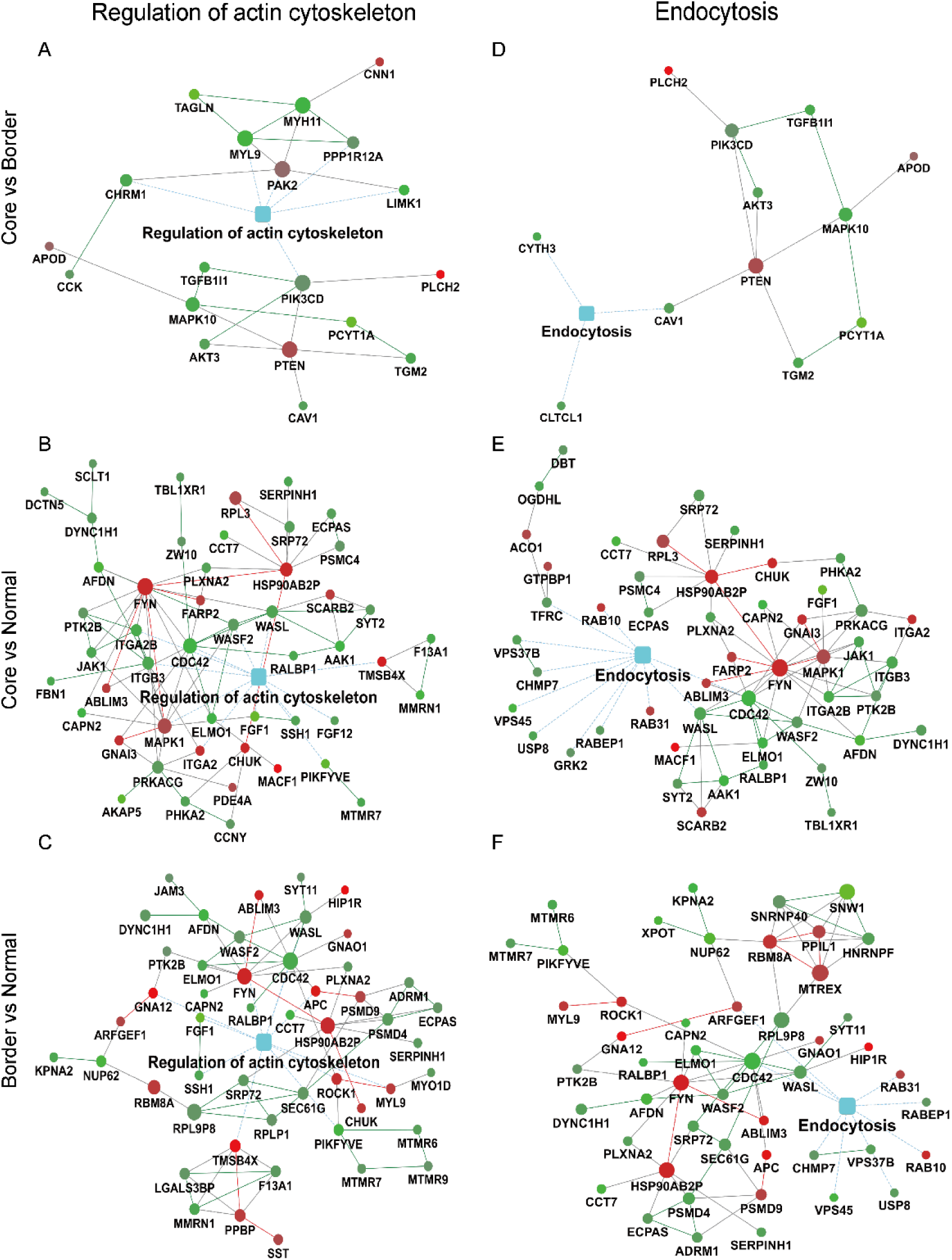
Protein-protein interaction (PPI) analysis of the DEPs involved in two common KEGG pathways for three group pairs. A. Regulation of actin cytoskeleton pathway from Core *vs.* Border group pair. B. Endocytosis pathway from Core *vs.* Border group pair. C. Regulation of actin cytoskeleton pathway from Core *vs.* Nonepilepsy group pair. D. Endocytosis pathway from Core *vs.* Nonepilepsy group pair. E. Regulation of actin cytoskeleton pathway from Border *vs.* Nonepilepsy group pair. F. Endocytosis pathway from Border *vs.* Nonepilepsy group pair.

We conducted a comprehensive analysis of the 412 DEPs that showed significant differences between the epileptic Core/Border groups and the Nonepileptic control samples (Figure 4A). To gain further insights into the functional implications of these DEPs, we subjected them to GO ontology and KEGG pathway analysis.

**Figure 4.**
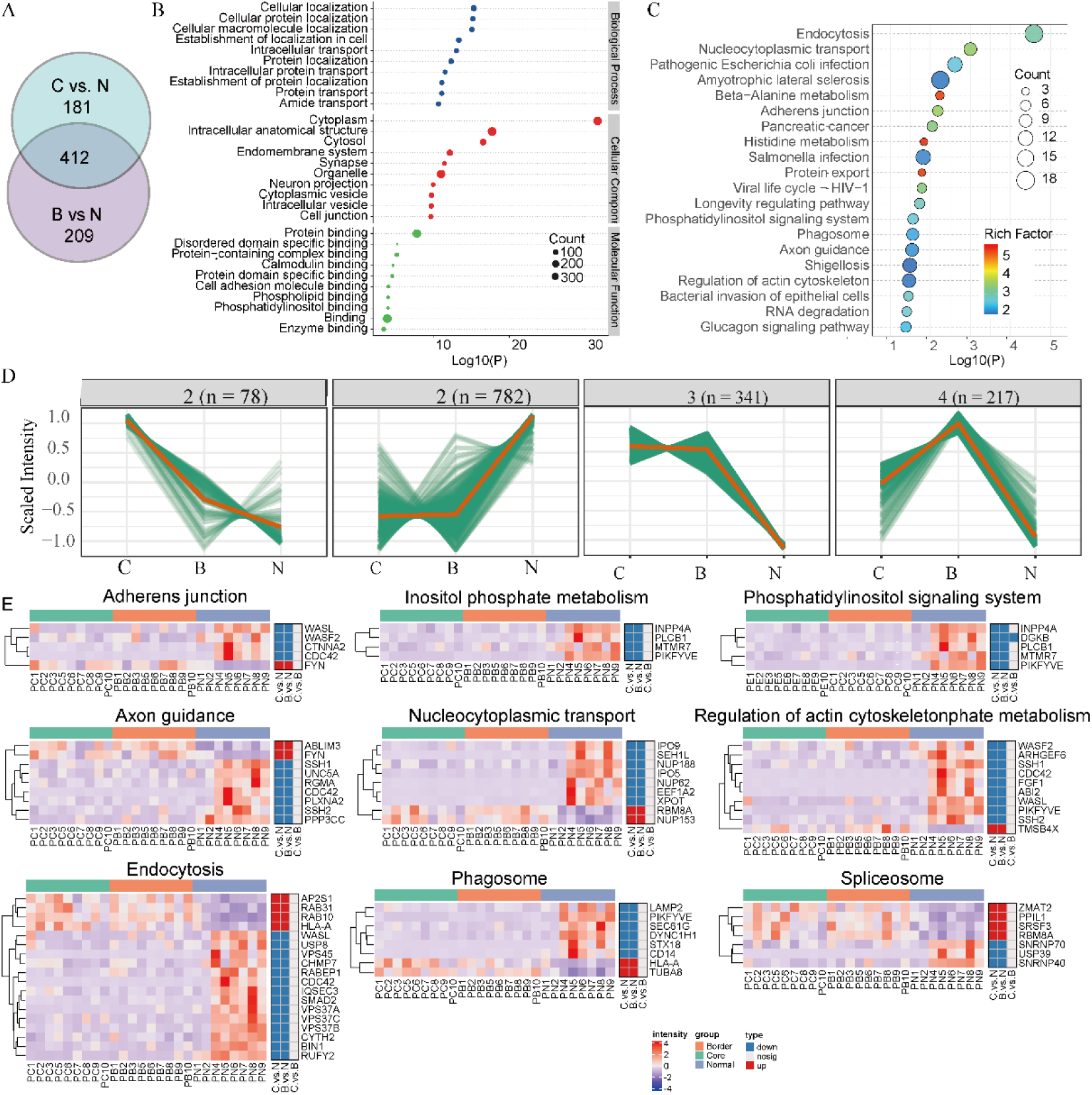
Analysis of DEPs between epilepsy and non-epilepsy patients. A. Venn statistics of DEPs of between epilepsy and non-epilepsy patients. B. GO analysis of the DEPs between epilepsy and non-epilepsy patients. C. KEGG analysis of the DEPs between epilepsy and non-epilepsy patients. D. K-means cluster analysis of DEPs between epilepsy and non-epilepsy patients. E. Cluster analysis of the DEPs of a few top different KEGG pathways between epilepsy and non-epilepsy patients.

In the GO analysis, we identified the top 10 terms for CC, MP, and BP, which are presented in Figure B. These terms provide valuable information about the cellular localization, molecular functions, and biological processes associated with the DEPs in epileptic patients.

Furthermore, the KEGG pathways analysis revealed the top 20 pathways based on their *p*-value and enrich factor (Figure 4 C). This analysis helped us identify the key pathways that are significantly altered in epileptic patients compared to non-epileptic individuals.

To explore the expression patterns of the DEPs among the three groups, we employed k-means cluster analysis with the K value determined according to the Calinski criterion (Figure 4 D). The majority of these trends (Figure 4 D 1 2 3) showed a clear separation between epileptic 4 and non-epileptic patients, indicating distinct expression patterns associated with epilepsy.

The major differences in pathways between epileptic and non-epileptic patients were found to involve various cellular processes, such as Endocytosis, Nucleocytoplasmic transport, Adherents junction, Spliceosome, Phosphatidylinositol signaling system, Phagosome, Axon guidance, and Regulation of actin cytoskeleton. Figure 4 E displays the main proteins involved in these pathways and their corresponding expression levels.

*Comparison of our DEPs with other proteomic data* To further validate the DEPs we identified, we searched for epilepsy proteomic data in the literature and found a substantial increase in the number of publications on this topic after 2020. The proteomic detection methods have evolved from the earlier two-dimensional electrophoresis to mass spectrometry, resulting in relatively varied results. However, the mass spectrometry-based detection methods provide more abundant results, so we mainly compared our data with literature published after 2020. We retrieved 70 literature reports using *epilepsy* and *proteomic* as keywords, and we found six literature reports with clear and reliable proteomics data that compared the epileptic group with a nonepileptic control group, however, none of them compared the differences between the core and border areas of epileptic foci. We summarized the data from these six literature reports in Table 3(Do et al, 2020; Pires et al, 2021; Xiao et al, 2020; Xu et al, 2021; Zhang et al, 2020; Zheng et al, 2020), and compared the DEPs identified in our study with the data from these reports, the statistic was shown in supplement Table Ev1. Form the statistic we see, the almost all of common protein molecules are included in our data and Geoffrey’s literature(28), so we further analyzed the common proteins those come with our data and Geoffrey’s literature and finally focused on 9 proteins: Q01995/TAGLN and P21810/BGN from C vs. B group pair, Q01469/FABP5, Q8TB36/GDAP1, O60888-3/CUTA and P46821/MAP1B from C vs. N and B vs. N group pairs, O95139/NDUFB6, Q2M2I8/AAK1 from C vs. N group pair, P30043/BLVRB from B vs. N group pair (see Table S2).

**Table 3:** Literatures on epileptic proteomic analyses had been compared with our study.

|  | Sample source | Samples of Lesions | Samples of control | Data sources in the literature | Results | Standard for differential protein | Detection method | Literature resources |
| --- | --- | --- | --- | --- | --- | --- | --- | --- |
| 1 | Rat Hippocampus | PTZ-kindled epileptic rats | equal dose of normal saline | Supplement table 3 | 6 downregulated, 93 upregulated | fold change >1.2 or <0.83, and p<.05 | iTRAQ labeling | Weiye Xu et,al(23). |
| 2 | Rat hippocampus | electric stimulation rodent model | sham control | Supplement table 1,4,7,10 | Supplement Table 1: 32 downregulated, 26 upregulated<br>Supplement Table 4: 11 downregulated, 5 upregulated<br>Tupplement Table 7: 52 downregulate, 22 upregulated<br>upplement Table 10: 29 downregulated, 5 upregulated | p<0.05 | Micro-proteomics | Amanda M. doCanto, et,al(24). |
| 3 | Rat hippocampus | lithium chloride-pilocarpine | 0.9% saline solution | Supplement table 1 | 92 downregulated, 18 upregulated | fold-change > 1.2 or < 0.83 and p < 0.05 | TMT labeling | Yu-qin Zheng et,al(25). |
| 4 | Human hippocampi | 4 TLE patients | autopsies of patients who died of non-neurological diseases | Supplement table 2c | 141 downregulated, 70 upregulated | >1.5-fold or <0.67-fold and p<.05 | TMT labeling | Yusheng Zhang et,al(26). |
| 5 | Human Hippocampi | 7 TLE patients | autopsied patients without disorders in the CNS | Supplement table 1 | 90 downregulated<br>82 upregulated | fold-change>2, or <0.5 and $p<0.01$ | iTRAQ-based | Wenbiao Xiao et,al(27). |
| 6 | Human Hippocampus DG | Autopsy epilepsy patients | Autopsy control patients | Supplement table 11 | 26 downregulated,<br>23 upregulated | Student's *t*-test, FDR <5% | label-free | Geoffrey Pires et,al(28). |
|  | Human Hippocampus CA1-3 |  |  | Supplement table 7 | 470 downregulated,<br>307 upregulated |  |  |  |
| 7 | Human Hippocampus or temporal lobe cortex | The core region of epileptogenic focus (C) | Margon of epileptogenic focus (B), Nonepileptic samples (N) | Datasets S1 and S2 | C vs B:<br>98 downregulated,<br>65 upregulated,<br>C vs N:<br>446 downregulated,<br>147 upregulated,<br>B vs N:<br>435 downregulated,<br>186 upregulated, | C vs B: fold change >1.2 or <0.83,<br>C vs N, B vs N: fold change>2 or <0.5, and $p<.05$, | DIA | this study |

*Altered KCNC2 expression and its role in epileptogenesis* Aberrant electrical activity is the core for epileptogenesis. To verify the significance of our proteomic study, we selected a DEP of Kv3.2, a potassium ion channel, for further study, which showed differential expression between Core *vs.* Border groups (FC _(C_ *_vs._* _B)_ = 0.235, p < 0.05). *KCNC2* encodes Kv3.2, a voltage-gated potassium channel of Kv3 subfamily which are essential for shaping communication and controlling excitability within the central nervous system (Lien & Jonas, 2003). Kv3.2 expression is almost limited to the brain, predominantly in the GABAergic interneurons in cortex and hippocampus (Rudy & McBain, 2001).

Firstly, we conducted immunohistochemistry to verify the expression of Kv3.2 in human temporal lobe epilepsy (TLE) tissue. The results demonstrated a significant decrease in the expression of Kv3.2 in the Core group compared to the Border group (Figure 5 A) (*p*<0.05), which is consistent with the proteomic results.

**Figure 5.**
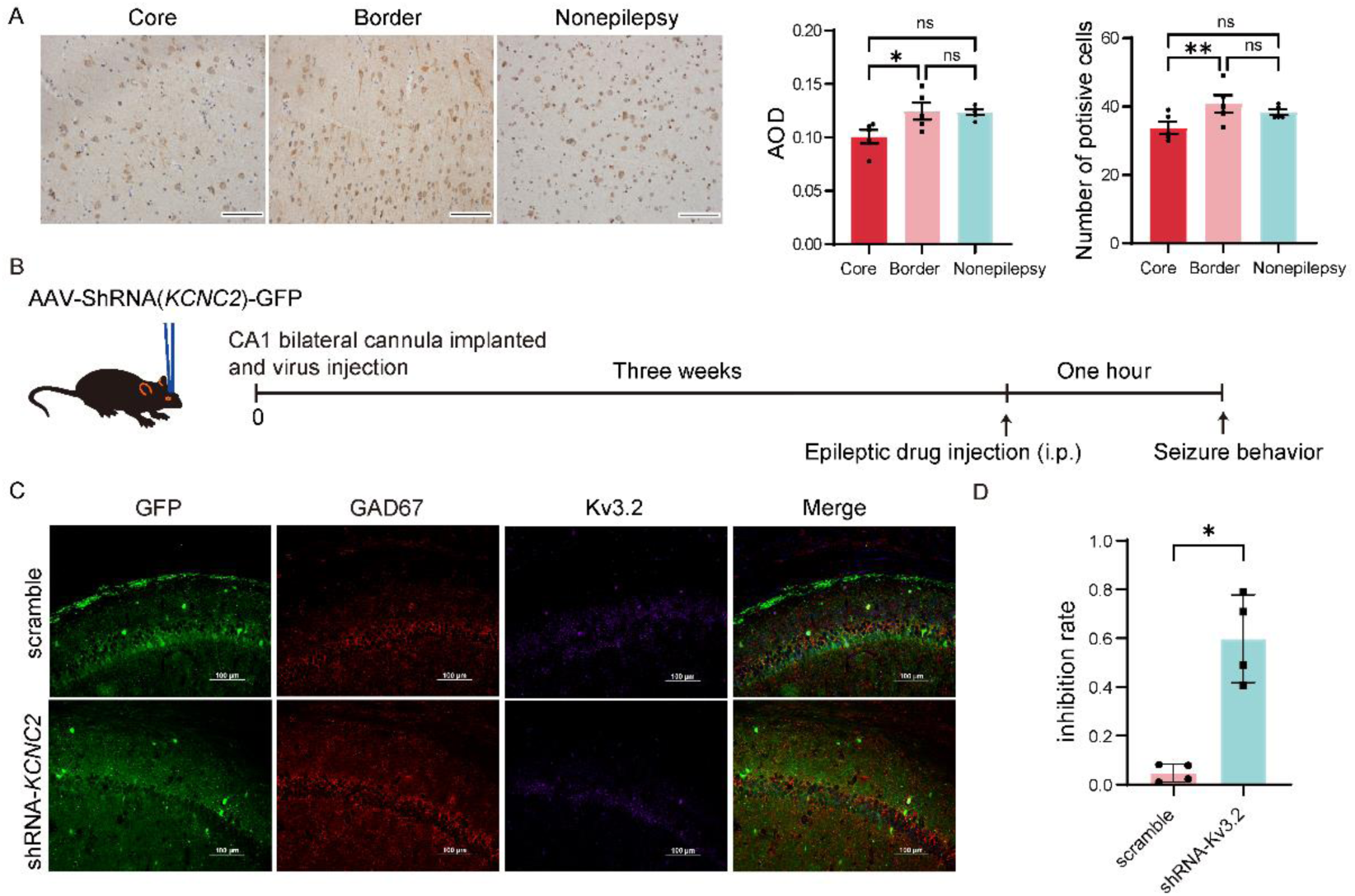
*KCNC2* expression in epileptic tissure. A. *KCNC2* expression and the statistics results of Core, Border and Nonepilepsy groups by immunohistochemical analysis in human tissue (n=5 for all three groups respectively, Data are presented as the mean ± SEM, AOD: Core 0.1008 ± 0.006368, Border 0.1247 ± 0.007888, Nonepilepsy 0.1235 ± 0.002691; positive cell number: Core 33.8 ± 1.772, Border 43.2 ± 1.985, Nonepilepsy 38.8 ± 0.7348). Design for study of Kv3.2 functions in mouse epileptic seizure model. B. Kv3.2 expression and its co-localization with GABAergic neurons in C57/6J mice infected with shRNA-*KCNC2* virus or scramble virus. C. The inhibition rate of shRNA-*KCNC2* virus on Kv3.2 expression in mice (mean ± SEM, scramble: 0.04793 ± 0.01849, shRNA-*KCNC2*: 0.5982 ± 0.08995, n=4)

To investigate the role of Kv3.2 in epileptic disorders, we conducted experiments using C57/6J mice. We injected a shRNA against *KCNC2* (AAV9-shRNA (*KCNC2*)-GFP) (shRNA-*KCNC2*) into the hippocampus (CA1) and confirmed the expression of the shRNA by observing the expression of GFP three weeks later (Figure 5 B and C). Compared to the scramble group (AAV9-scramble shRNA-GFP), the expression of Kv3.2 was significantly reduced in the shRNA-*KCNC2* group, as measured by qPCR (Figure 5 D); Immunofluorescence studies further confirmed the reduction of Kv3.2 in GABAergic neurons within the hippocampus of the mice (Figure 5 C).

Next, we established four different epileptic models in mice using intraperitoneal injections of pilocarpine, pentylenetetrazol (PTZ), penicillin, and hymen acid (KA), respectively. We then observed seizures in the mice. The results showed that the latency of grade 3 seizures in the shRNA-*KCNC2* group was significantly shorter than that in the scramble group for all four models (independent sample *t*-test: pilocarpine, PTZ, KA, *p*<0.01; penicillin *p*<0.05) (Figure 6 A). Additionally, the incidence of grade 3, grade 4, grade 5, and above seizures increased significantly in the pilocarpine and KA models in the shRNA-*KCNC2* group compared to the scramble group (chi-square test: pilocarpine *p*<0.001, KA grade 4 and above *p*<0.001, KA grade 3, grade 5 and above *p*<0.05). In the PTZ and penicillin models, the incidence of grade 4 and above seizures increased significantly (*p*<0.05) (Figure 6 B). These findings suggest that reduced expression of Kv3.2 increases susceptibility to epilepsy, and decreased expression of Kv3.2 may be a significant factor contributing to epileptic seizures. Notably, the pilocarpine model emerged as the most suitable animal model for this study.

**Figure 6.**
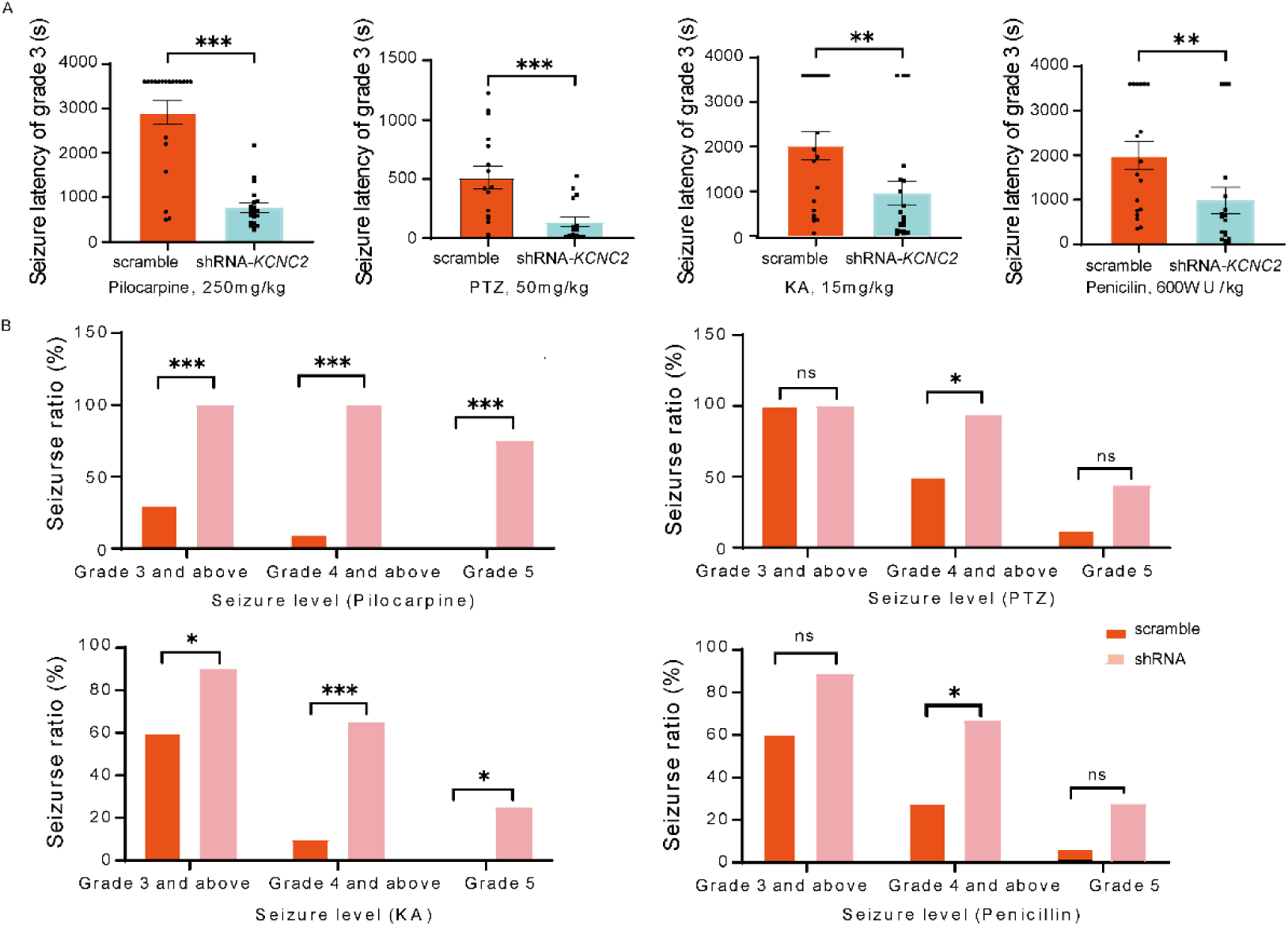
The action of *KCNC2* in mouse epileptic seizure model. A. The statistics of seizure latency of grade 3 in the mouse seizure model induced with KA, PTZ, Pilocarpine and Penicillin, respectively. Data are presented as the mean±SEM, Pilocarpine: scramble 2911±260.2, shRNA-*KCNC2* 767.9±102.9, n=20 for both groups; PTZ scramble 512.1±96.94, shRNA-*KCNC2* 139.4±42.65, n=16 for both groups; KA scramble 2911±260.2, shRNA-*KCNC2* 767.9±102.9, n=20 for both groups; Penicillin scramble 1995±311, shRNA-*KCNC2* 989.1±297.5, n=18 for both groups. B. The statistics of seizure ratio of the mouse seizure model induced with KA, PTZ, Pilocarpine and Penicillin, respectively. Pilocarpine grade 3 and above scramble 30.00%, shRNA-*KCNC2* 100%, grade 4 and above scramble 10.00%, shRNA-*KCNC2* 100%, grade 5 and above scramble 0, shRNA-*KCNC2* 75.00%(n=20); PTZ grade 3 and above scramble 100%, shRNA-*KCNC2* 100%, grade 4 and above scramble 50.00%, shRNA-*KCNC2* 93.75%, grade 5 and above scramble 12.50%, shRNA-*KCNC2* 43.75% (n=16); KA grade 3 and above scramble 60.00%, shRNA-*KCNC2* 90.00%, grade 4 and above scramble 10.00%, shRNA-*KCNC2* 65.00%, grade 5 and above scramble 0, shRNA-*KCNC2* 25.00% (n=20); Penicillin grade 3 and above scramble 61.11%, shRNA-*KCNC2* 88.89%, grade 4 and above scramble 27.78%, shRNA-*KCNC2* 66.67%, grade 5 and above scramble 5.56%, shRNA-*KCNC2* 27.78% (n=18). Note: ns (*p*>0.05), *(*p*<0.05), **(*p*<0.01), ***(*p*<0.001).

*Mechanism for Kv3.2 involvement in epileptogenesis* The role of Kv3.2 in electrical discharges of neurons in the CA1 region of the mouse hippocampus was investigated using whole-cell patch clamp recordings. Brain slices were obtained from mice previously injected (three weeks) with either shRNA-*KCNC2* or scramble shRNA (Figure 7 A). Action potentials (APs) were induced by a ramp or a step current clamp, and firing frequency of APs and the stimulation intensity required for the occurrence of APs (rheobase) were recorded. To determine the identity of the recorded neurons, single-cell PCR was performed after the electrophysiological recordings (some of single-cell PCR results shown in Figure 7 B).

**Figure 7.**
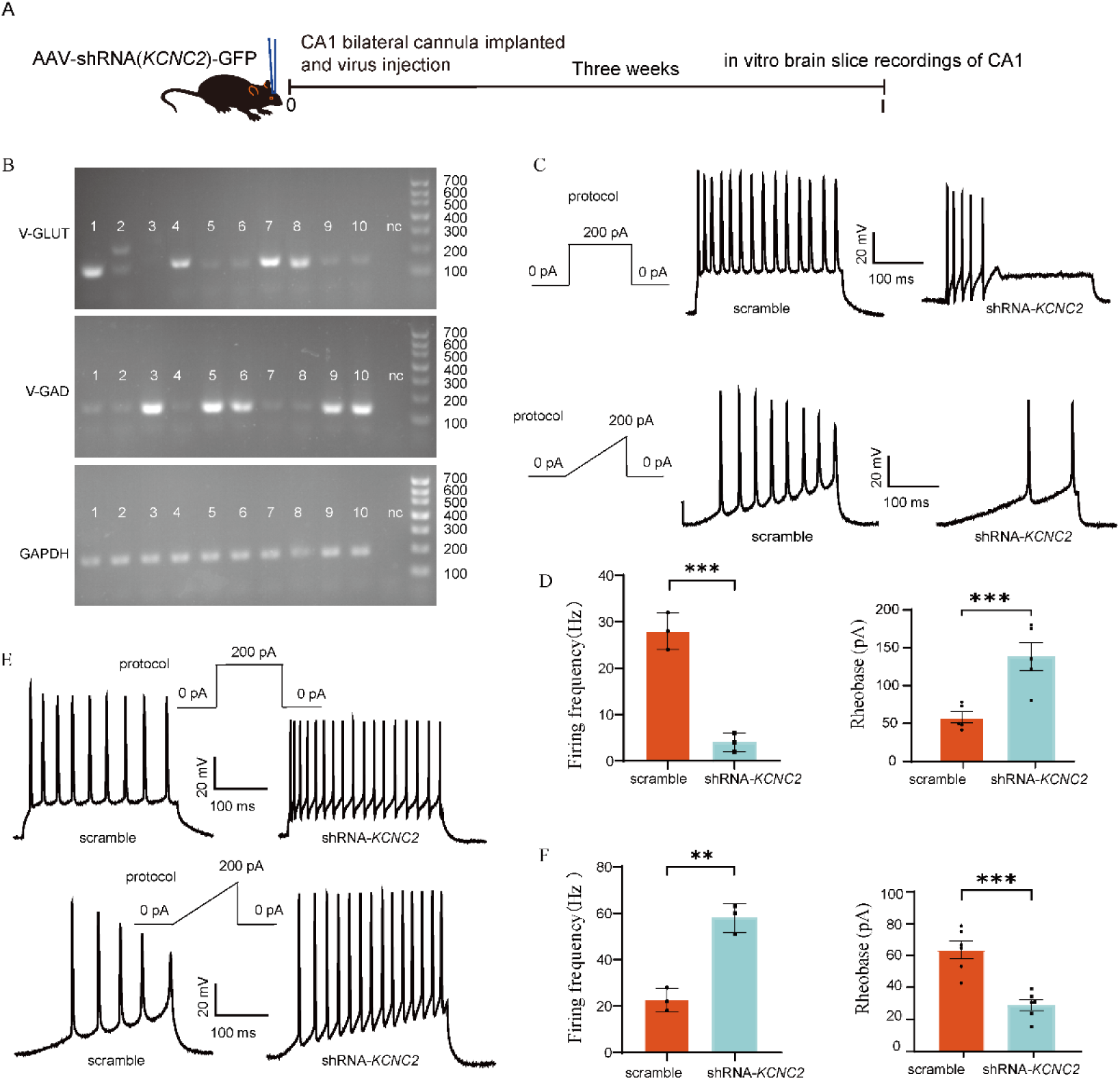
Mechanism for Kv3.2 involvement in initiation of epileptic electrical firing with mouse model. A. The study design for mechanism of Kv3.2 in epileptic seizure. B. Some of the single cell PCR results. C. Firing frequency and cell excitability of GABA neurons in brain slices which from C57/6J mice infected with shRNA-*KCNC2* virus or scramble virus. D. The statistics result of firing frequency and rheobase of GABA neurons (Data are presented as the mean±SD, rheobase: scramble 58.26±7.346, shRNA-*KCNC2* 138.6±18.34, n=5 for both groups; firing frequency scramble 28±2.309, shRNA-*KCNC2* 4±1.155, n=3 for both groups). E. Discharge frequency and cell excitability of glutaminergic neurons in brain slices which from C57/6J mice infected with shRNA-*KCNC2* virus or scramble virus. F. The statistics result of firing frequency and rheobase of glutaminergic neurons (Data are presented as the mean±SD, rheobase, scramble 63.48±5.53, shRNA-*KCNC2* 28.97±3.417, n=5 for both groups; firing frequency, scramble 22.67±2.906, shRNA-*KCNC2* 58±3.606, n=3 for both groups). Note: **(*p*<0.01), ***(*p*<0.001).

The results indicated that knocking down Kv3.2 reduced the firing frequency of APs and increased the Rheobase in GABAergic neurons (Figure 7 C and D), while the opposite effects were observed in glutaminergic neurons (Figure 7 E and F).

To further investigate the correlation between Kv3.2 and epilepsy, cortical and intracranial electrodes were placed in C57/6J mice to record EEG and analyze the discharge characteristics of the hippocampal CA1 region (Figure 8 A). The EEG results recorded by intracranial electrodes (Figure 8 B) showed that, compared to the scramble group and wild type group, the amplitude and discharge frequency are much higher, the power density curve shifted upward and the power increased in the shRNA-*KCNC2* group, including the power of full spectrum band and α, β and γ rhythm (Figure 8 C). Time-frequency analysis and total power statistical results also revealed a similar pattern of changes (Figure 8 D and E). Highly consistent results were also obtained from cortical EEG recordings (Figure 9 A and B).

**Figure 8.**
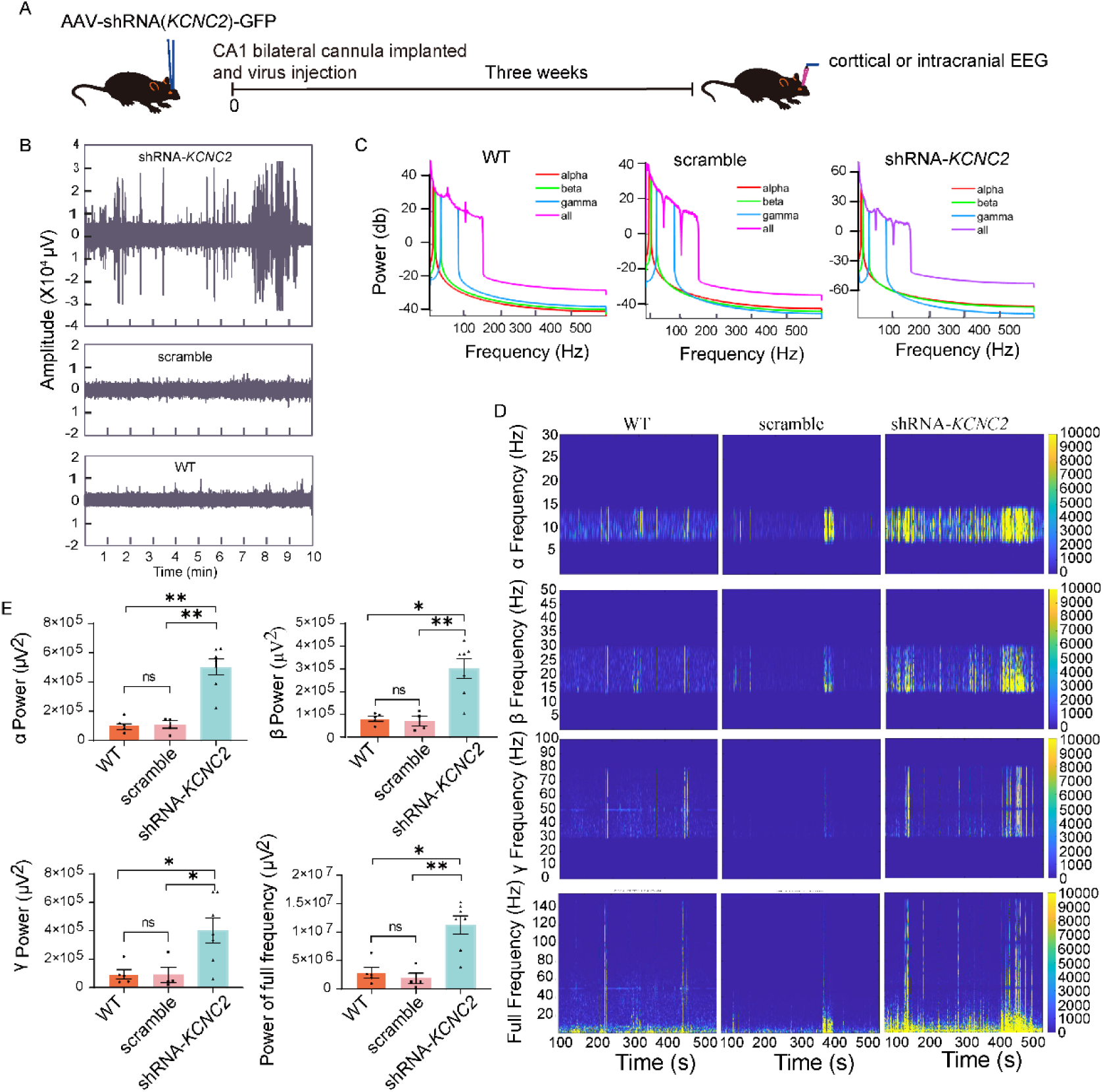
The action of altered Kv3.2 expression on mouse intracranial EEG. A. Study design. B. A typical example for intracranial EEG recording of wild type C57/6J mice and mice infected with shRNA-*KCNC2* virus or scramble virus. C. The power density curve of α, β, γ rhythm and full spectrum band for panel B. D. Time-frequency analysis α, β, γ rhythm and full spectrum band for panel B. E. The statistics result of the power of α, β, γ rhythm and full spectrum band. Data are presented as the mean ± SEM, α frequency, wild type 104869 ± 21122, scramble 102796 ± 26834, shRNA-*KCNC2* 499236 ± 55353; β frequency, wild type 81961 ± 10320, scramble 68932 ± 20292, shRNA-*KCNC2* 300537 ± 43759; γ frequency, wild type 94387 ± 32402, scramble 89761 ± 53239, shRNA-*KCNC2* 403460 ± 87842; full frequency, wild type 2872886 ± 923100, scramble 1840753 ± 927895, shRNA-*KCNC2* 11263651 ± 1612478. shRNA-*KCNC2* n=5, scramble n=5 wild type n=3 and 16 recording channels for each sample). Note: ns (*p*>0.05), *(*p*<0.05), **(*p*<0.01), ***(*p*<0.001).

**Figure 9.**
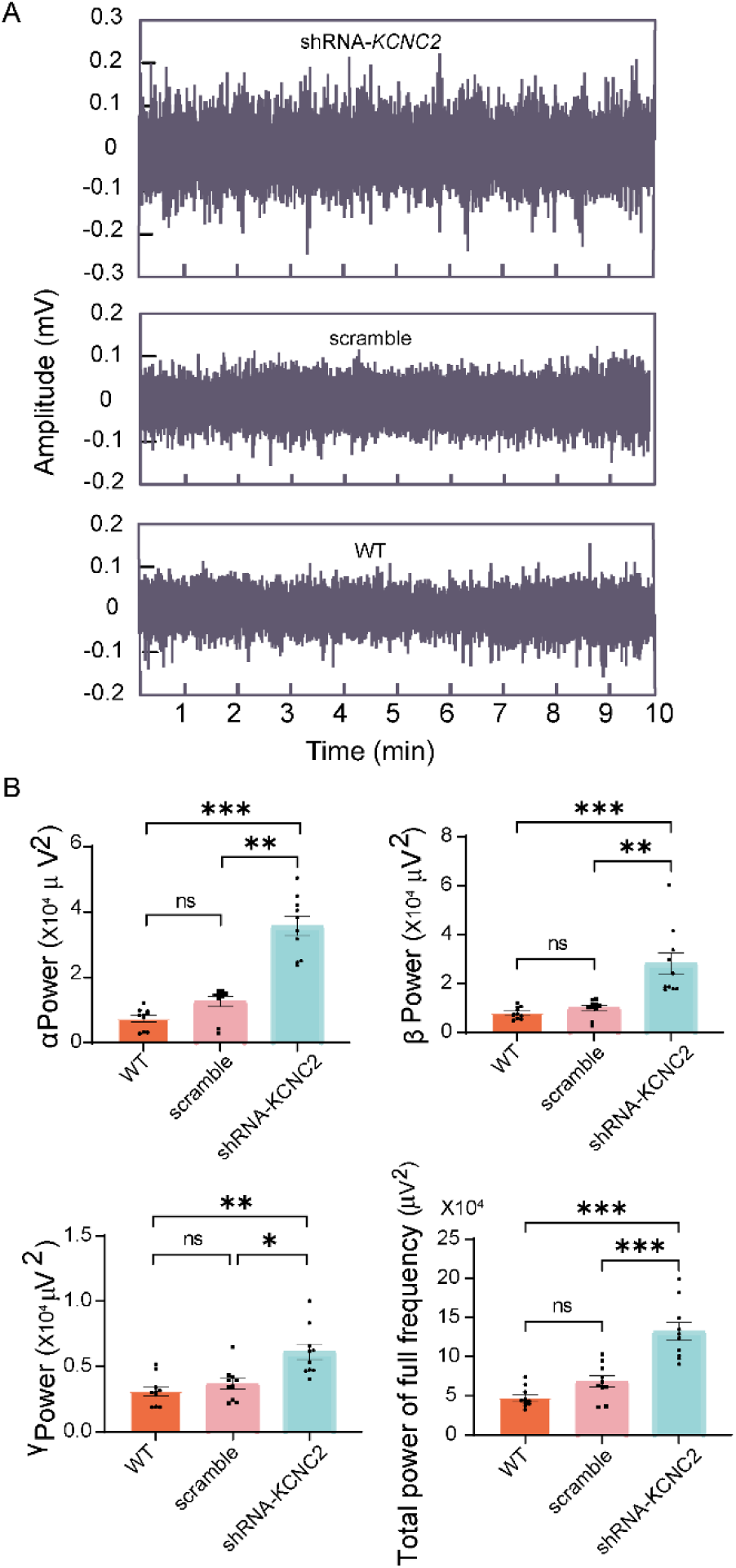
The action of altered Kv3.2 expression on mouse cortical EEG. A. A typical example for cortical EEG recording of wild type C57/6J mice and mice infected with shRNA-*KCNC2* virus or scramble virus. B. The statistics result of total power of α, β, γ rhythm and full spectrum band for cortical EEG. Data are presented as the mean ± SEM, α frequency, wild type 0.7423 ± 0.1029, scramble 1.26 ± 0.1515, shRNA-*KCNC2* 3.572 ± 0.2929, β frequency, wild type 0.8006 ± 0.07738, scramble 0.9878 ± 0.1117, shRNA-*KCNC2* 2.815 ± 0.4439; γ frequency, wild type 0.3108 ± 0.03605, scramble 0.3661 ± 0.0406, shRNA-*KCNC2* 0.6112 ± 0.05895; full frequency, wild type 4.769 ± 0.409, scramble 6.82 ± 0.7013, shRNA-*KCNC2* 13.2 ± 1.145. shRNA-*KCNC2* n=7, scramble n=4 wild type n=5. Note: ns (*p*>0.05), *(*p*<0.05), **(*p*<0.01), ***(*p*<0.001).

These findings demonstrate the significant role of Kv3.2 in modulating neuronal excitability in the CA1 region of the hippocampus. Knocking down Kv3.2 led to altered firing frequencies and Rheobase in GABAergic and glutaminergic neurons, suggesting its influence on inhibitory and excitatory signaling in the brain. The observed changes in EEG power density and time-frequency analysis further support the correlation between Kv3.2 and epileptic activity in the CA1 region.

## Discussion

In this study, we conducted an extensive proteomic analysis using human surgical samples obtained from both epileptic and non-epileptic patients. Our primary focus was to investigate the functional roles of differentially expressed proteins (DEPs) in epileptogenesis. Notably, we not only compared the proteomics analysis and function evaluation of DEPs between epileptic and non-epileptic patients but also compared these aspects between samples from the core and the border regions of the epileptic focus. The investigation of these distinct regions is crucial as they may play different roles in the process of epileptogenesis. Our study yielded multiple novel findings that provide valuable insights into the mechanism of epileptogenesis. These discoveries are expected to inspire new perspectives and thoughts in the field.

Prior to our study, proteomic analyses had been conducted on both rat epileptic models and human epileptic samples. While some literature reported comparative studies between epileptic patients and autopsy samples from deceased individuals, none had specifically explored the differences between the core and the border regions of the epileptic focus within the same patients (Table 3). One key observation from these literatures was that there was little similarity in the DEPs between rat models and human samples. Furthermore, the results obtained from human samples varied significantly across different studies, likely due to various factors such as sample sources, detection methods, and screening criteria. In particular, the choice of detection method and the number of differential proteins screened were found to have a significant impact on these variations. Through the comparisons, we got 9 common proteins at least in 3 studies. In our study, we employed the DIA proteomic detection method, which was chosen to overcome the limitations of label-free or isobaric tags technology. The DIA method involves the systematic collection of MS2 spectra, covering the entire MS1 spectrum, ensuring comprehensive and accurate protein identification(Pappireddi et al, 2019). To establish an appropriate nonepileptic control group, we utilized brain tissue surgically removed from non-epileptic patients with conditions such as trauma and cerebrovascular accidents. We also included non-tumor patients to exclude any interference from tumor-related pathogenic factors. For our case control group, we used the distal lesion tissue from the same epileptic patient. This approach helped minimize the influence of various interfering factors and provided the most effective protein molecular library for analysis. Our study aimed to investigate both differentially expressed proteins (DEPs) between the core and border areas of the epileptogenic focus and DEPs between epileptic patients and non-epileptic patients. This rigorous approach enabled us to more precisely screen and identify key protein molecules associated with epileptic seizures and epileptogenic pathogenesis. By comparing these distinct groups, we aimed to gain deeper insights into the underlying mechanisms of epilepsy.

One of the most intriguing findings of our study is related to the 10 differentially expressed proteins (DEPs) that exhibited consistent alterations between two pairs of groups: the Core *vs.* Border group and the Core *vs.* Nonepileptic control group. Among these DEPs, 7 were expressed at lower levels in the Core group compared to both the Border and Nonepileptic control groups. Conversely, 3 DEPs were expressed at higher levels in the Core group compared to both the Border and Nonepileptic control groups. This consistency in expression patterns indicates the potential significance of these proteins in distinguishing the epileptic core from the border region and the non-epileptic control group, indicating a possible role in epileptogenesis.

The three DEPs mentioned earlier which expressed at higher levels in the Core group — P35754/GLRX, O75335/PPFIA4, and Q96KP4/CNDP2—hold significant implications within our study. GLRX is characterized by its glutathione-disulfide oxidoreductase activity, primarily involved in the reduction of low molecular weight disulfides and proteins susceptible to glutathionylation. This action can modulate several functional pathways including cytoskeletal dynamics, glycolysis/energy metabolism, kinase-mediated signaling, calcium homeostasis, antioxidant defense, and protein folding. Despite its relevance, only limited evidence connects GLRX to epilepsy, warranting further exploration(A et al, 2022; Ogata et al, 2021).

PPFIA4, on the other hand, is associated with the regulation of focal adhesion disassembly and the precise localization of receptor-like tyrosine phosphatases type 2A on the plasma membrane. This regulation may influence interactions with the extracellular environment and substrate associations. While PPFIA4’s role has been observed predominantly in various cancer contexts, there’s been little exploration of its relation to epilepsy, barring a case report involving drug-resistant focal onset epilepsy in connection with PPFIA4 gene variants(Sourbron et al, 2021; Xie et al, 2020; Xu et al, 2022; Zhao et al, 2022; Zhou et al, 2022).

As for CNDP2, the human carnosine dipeptidase, it exists in two isomeric forms—CNDP1 and CNDP2—distinguished by their activators (Zn^2+^ in serum for CNDP1 and Mn^2+^ in tissue for CNDP2). CNDP2 exhibits hydrolytic activity on various dipeptides and can activate the mitogen-activated protein kinase (MAPK) pathway, thereby promoting cell proliferation. While imbalances in CNDP2 activity and expression have been linked to multiple diseases, including neurological conditions, recent research proposes CNDP levels as potential markers for neurological diseases. Nonetheless, the association between CNDP and epilepsy remains unexplored (1983; Zhang et al, 2014).

Within our study, we have highlighted a particularly significant discovery involving the potassium channel Kv3.2, which is encoded by *KCNC2*. Notably, the expression of Kv3.2 was found to be reduced in the Core group. The Kv3 subfamily encompasses four members: Kv3.1, Kv3.2, Kv3.3, and Kv3.4. These channels exhibit unique characteristics, opening at a relatively depolarized membrane potential of −10 mV and displaying rapid deactivation rates approximately 10 times faster than other potassium channels(Kaczmarek & Zhang, 2017).

Kv3.2 finds its predominant expression within the brain, specifically in GABAergic interneurons situated in the cortex and hippocampus(Rudy & McBain, 2001). Its role is indispensable for regulating communication and excitability within the central nervous system, crucial for maintaining a balanced neural environment(Lien & Jonas, 2003). Altered Kv3.2 function can lead to hyperexcitability in GABAergic interneurons, a condition implicated in epilepsy(Erisir et al, 1999).

In the context of epilepsy, the significance of Kv3.2 gains further support from patient cases. A de novo variant in *KCNC2* was identified in a patient with NF1-WS (Neurofibromatosis type 1 with Legius syndrome) and another with epileptic encephalopathies, firmly establishing the role of Kv3.2 in epilepsy(Rademacher et al, 2020; Vetri et al, 2020). Subsequent research has reported *KCNC2* mutations in additional cases of epilepsy(Rydzanicz et al, 2021).

Our study’s findings, combined with these previous observations, bolster the notion that Kv3.2 plays a pivotal role in epileptogenesis. This involvement is characterized by its impact on GABAergic neuron activity and ultimately contributes to our understanding of the mechanisms underlying epilepsy.

In conclusion, our comprehensive proteomic analysis of human surgical samples from both epileptic and non-epileptic patients, with a specific focus on DEPs, has yielded invaluable insights into the complex process of epileptogenesis. By exploring DEPs between epileptic patients and non-epileptic patients, as well as between the core and border regions of the epileptic focus, we have unveiled novel findings that shed light on the underlying mechanisms of epilepsy. We hope that these findings will serve as a catalyst for future research, inspiring new avenues of investigation and deepening our comprehension of epileptogenesis. Ultimately, our study contributes to the growing body of knowledge aimed at developing more targeted therapeutic strategies for individuals affected by epilepsy.

## Materials and Methods

### Study design

In this study, our objective is to identify key molecules involved in epileptogenesis. To achieve this, we utilized proteomic data obtained from specimens collected from 9 epileptic patients and 8 non-epileptic patients. These specimens were categorized into three distinct groups: the core regions of focal tissue from epileptic patients (referred to as the Core group), the border regions of focal tissue from the same epileptic patients (referred to as the Border group), and a control group consisting of non-epileptic patients (referred to as the Nonepilepsy group). We employed Data-Independent Acquisition (DIA) methods to analyze these samples.

Differentially expressed proteins and peptides (DEPs) between the Core and Border groups were identified as a molecular pool potentially associated with aberrant electrical discharges. Additionally, DEPs between epileptic patients and non-epileptic patients were considered as a molecular pool relevant to epileptogenesis. From these pools, we selected a specific protein, Kv3.2, for further investigation. To assess the role of Kv3.2, we designed AAV9-shRNA-*KCNC2* and scramble virus constructs, which were subsequently injected into the mouse CA1 region of the hippocampus. After a 3-week period of the virus injection, we induced an epilepsy model to evaluate epilepsy susceptibility and seizure activity. Cortical electroencephalogram (EEG) and intracranial electroencephalogram recordings were employed to monitor EEG patterns.

Additionally, we utilized brain patch recordings to monitor cell firing and elucidate the mechanism underlying the action of Kv3.2 in epilepsy seizure. The current study conforms with World Medical Association Declaration of Helsinki and the Guide for the Care and Use of Laboratory Animals from National Natural Science Foundation of China. The work was approved by the Medical Ethics Committee of Hebei Medical University and undertaken with the understanding and written consent of each subject (No. 2020-R240; No. 2022057)

### Samples and group

The excised tissue was from 9 cases of refractory medial temporal lobe epilepsy patients diagnosed by the neurosurgery department of the Second Hospital of Hebei Province. The information of patients was shown in table 1. The epileptic focus was determined by stereotactic electroencephalogram and electrophysiological monitoring during the operation. Out of the 9 patients, 8 underwent anterior temporal lobectomy, which involves the amygdala, hippocampus, and parahippocampal gyrus. Additionally, one patient underwent resection of the right anterior temporal lobe and the right temporal base based on the findings from the stereotactic electroencephalogram. The excised tissue of control group was collected from 8 patients with non-epileptic disease who undergoing emergency resection in the Department of Neurosurgery of Hebei Medical University. Inclusion criteria: (1) no history of neurological diseases; (2) no history of epilepsy and family history; (3) temporal lobe resection; (4) no history of glioma and other tumors. The brain tissue was removed according to the predetermined operation. The brain tissues were cut into small pieces about 1cm^3^ and immediately transferred to the frozen storage tube and stored in a -80°C refrigerator. All operations completed within 5 minutes. Specimen collection was approved by the ethics committee of the hospital and all patients signed informed consent.

### DIA proteomic experiment

*Reagents* Ammonium bicarbonate, dithiothreitol (DTT), iodoacetamide (IAA), and sodium carbonate were purchased from Sigma-Aldrich (St. Louis, MO). Urea and Sodium dodecyl sulfate (SDS) were purchased from Bio-Rad (Hercules, CA). Acetonitrile and water for nano-LC−MS/MS were purchased from J. T. Baker (Phillipsburg, NJ). Trypsin was purchased from Promega (Madison, WI). All other chemical reagents were purchased with analytical grade.

*Sample preparation* Protein was extracted from tissue samples using SDT lysis buffer and quantified with a BCA Protein Assay Kit (BeyoTime, China). Protein digestion was performed with FASP method described by Wisniewski, Zougmanet al.(Wisniewski et al, 2009). The concentrations of re-dissolved peptides were determined with OD280 by Nanodrop One device (Thermo, USA). Pooled peptide mixture of each sample was fractionated using a WatersXBridge BEH130 column (C18, 3.5 μm, 2.1×150 mm) on an Agilent1260 high-performance liquid chromatographer (HPLC) operating at 0.3 mL/min. A total of 60 fractions were collected for each peptide mixture and pooled to 20 fractions for further LC-MS analysis.

*LC-MS/MS analysis for DDA* Peptides from each pooled fraction were first spiked with iRT standard peptides (Biognosys AG, Switzerland) and separated using reverse-phase high-performance liquid chromatography (RP-HPLC) on EASY-nLC system (Thermo Fisher Scientific, Bremen, Germany) with a column (75 μm × 150 mm; 2μm ReproSil-PurC18 beads, 120 Å, Dr. Maisch GmbH, A mmerbuch, Germany) at a flow rate of 300 nL/min. The eluted peptides were analyzed on a Q Exactive HF-X mass spectrometer. MS data was acquired using a data-dependent top20 method dynamically choosing the most abundant precursor ions from the survey scan (350–1500m/z) for HCD fragmentation. The instrument was run with peptide recognition mode enabled. A lock mass of 445.120025 Da was used as internal standard for mass calibration. The full MS scans were acquired at a resolution of 60,000 at m/z 200, and 15,000 at m/z 200 for MS/MS scan. The maximum injection time was set to 50 ms for MS and 30ms for MS/MS respectively. Normalized collision energy was 28 and the isolation window was set to 1.6 Th. Dynamic exclusion duration was 30s.

*LC-MS/MS analysis for DIA* Peptides from each sample were spiked with iRT equally and separately. LC-MS/MS were also performed on a Q ExactiveHF-X mass spectrometer coupled with Easy 1200nLC (Thermo Fisher Scientific). The LC condition was set as the same as DDA method above. The DIA method consisted of a survey scan from 400-1200 m/zat resolution 60000 with AGC target of 3E6 and 30ms injection time. The 20 DIA MS/MS were acquired at resolution 15000 with 20 m/z isolation window and with AGC target of 1E6 and 50ms injection time. The Normalized collision energy was set 30. The spectra of full MS scan and DIA scan were recorded in profile and centroid type respectively.

*Sequence database searching* The DDA MS data were analyzed using MaxQuant software version 1.6.0.16. MS data were searched against the UniProtKB human database (186532 total entries, downloaded 10/2019) spiked in protein consisted with 11 iRT peptide sequences. The trypsin was selected as digestion enzyme. The maximal two missed cleavage sites and the mass tolerance of 4.5 ppm for precursor ions and 20ppm for fragment ions were defined for database search. Carbamidomethylation of cysteines was defined as fixed modification, while acetylation of protein N-terminal, oxidation of Methionine was set as variable modifications for database searching. The database search results were filtered and exported with <1% false discovery rate (FDR) at peptide-spectrum-matched level, and protein level, respectively.

*DIA raw data analysis* The DIA MS data were analyzed with Spectronaut pulsar X (Biognosys AG, Switzerland)(Barkovits et al, 2020). The Spectronaut was used for spectral library generation from Maxquant search results. The Spectronaut search were set as the default settings and the dynamic iRT was used for retention time prediction. The interference correction for MS/MS scan was enabled. The results were exported with <1% FDR at peptide level and protein level respectively.

*Proteome bioinformatics data analysis* Analyses of bioinformatics data were carried out with Perseus software(Tyanova et al, 2016), Microsoft Excel and R statistical computing software. Expression data were grouped together by hierarchical clustering according to the heatmap package, which is based on the open-source statistical language R25, using Euclidean distance as the distance metric and complete method as the agglomeration method. To annotate the sequences, information was extracted from UniProtKB or SwissProt database(Boutet et al, 2016), Kyoto Encyclopedia of Genes and Genomes (KEGG)(Kanehisa et al, 2012), and Gene Ontology (GO)(Ashburner et al, 2000). The GO terms were divided into three categories: biological process (BP), molecular function (MF), and cellular component (CC). GO and KEGG enrichment analyses were carried out with the Fisher’s exact test, and Benjamini–Hochberg false discovery rate (BH-FDR) correction for multiple tests was obtained. Construction of protein–protein interaction (PPI) networks were conducted by using the STRING database with the Cytoscape software(Kohl et al, 2011).

### Immunohistochemistry of human brain tissue

Fresh brain tissue was fixed with 4% paraformaldehyde and modified, dehydrated with gradient alcohol, embedded in paraffin, sliced with 10μm thick, and baked in oven at 60°C, antigen repaired, 3% hydrogen peroxide at room temperature and light protection for 20 min, 0.5 % triton membrane penetration, 10 % donkey serum closed at 37 °C for 30 min, primary antibody (anti-Kv3.2 antibody, 1:500, Cat.no. APC-011, alomone lab, Israel) was incubated at 4°C overnight. After rinsing with PBS, enzyme-labeled secondary antibody (1:200, Cat.no. A21206, Invitrogen, USA) incubated for 2 hours and visualized with DAB (Cat.no.D405772-1Set, Aladdin, China). The sections were observed and captured with a panoramic fluorescence machine the BX63 microimaging system (panoramic SCAN, 3D HISTECH, Hungary). For each group, five fields were randomly selected under 20 × magnification, Image Pro Plus 6.0 software was used to analyze the average optical density value and the number of positive cells of each reactive protein in different groups. Graphpad prism 9 software was used for statistical analysis.

### Immunofluorescence of C57 mice

The mice were intraperitoneally injected with pentobarbital (50 mg/kg) and then decapitated. the brain tissue was quickly removed and fixed in 4% PFA overnight at 4 °C after washed. Briefly, brain tissue was fixed, embedded, sliced, cleaned, sealed, and incubated in the primary antibody (anti-Kv3.2 antibody, 1:200, Cat.no. APC-011, alomone lab, Israel; anti-GAD67 antibody, 1:200, Cat.no. MAB5406, Sigma, USA) at 4°C overnight. After rinsed, slices were incubated with secondary antibody (1:200, Cat.no. A21206, Invitrogen, USA; Cat.no.A10037, Invitrogen, USA; 1:200, Cat.no.MM63010, Maokang Biology Co., Ltd, Shanghai, China) for 2 h at room temperature in dark and attached onto slides. The images were captured with the BX63 microimaging system (panoramic SCAN, 3D HISTECH, Hungary). Five fields were randomly selected under 20 × magnification for each group, Image Pro Plus 6.0 software was used for analysis and GraphPad prism 9 software was used for statistics.

Construction of shRNA AAV-9 Virus and Stereotactic Injection into Mouse Hippocampus Three pairs of siRNA sequences of the target genes were designed and transfected into Hek-293A cells. The siRNA with the highest silencing efficiency was screened by qPCR and packaged into shRNA AAV9 virus with GFP tags by Shandong Weizhen Biotechnology Co., LTD. Mouse *KCNC2* Gene ID: 268345, titer: 6.3×10^13^ vg/ml, shRNA sequence: GCAAGACAGAATTGAACATTTCAAGAGAATGTTCAATTCT GTCTTGCTTTTTT. C57/6J mice were anesthetized with 1% pentobarbital sodium (50mg/kg) and fixed in the stereotactic frame. After drilling the hole into the skull surface, 300 nL virus was bilaterally injected into hippocampus CA1 (AP: -2.50 mm; ML: ±2.0 mm; DV: -1.5mm) using an infusion pump (RWD Life Science, Shenzhen, China) at the rate 150 nL/min. 21 days after injection, the brain was exposed and the hippocampus was removed. qPCR was used to verify the knocking efficiency. After cardiac perfusion and prefixation with paraformaldehyde, hippocampus was taken, frozen section was taken, and the virus injection site was observed by fluorescence microscope. When Mice successfully injected and knocked down the target genes the mice used for subsequent study.

### qPCR

Total RNA was extracted from tissues or cells and reverse transcription into cDNA. qPCR reaction system: SYBR Premix Ex Taq II (2×) 12.5 ul, Forward Primer (10 uM) 1ul, Reverse Primer (10uM) 1ul, cDNA 2 ul, DEPC water 8.5 ul. Reaction condition: 1 cycle of 95°C for 0:30, followed by 39 cycles of 95°C for 0:05, 60°C for 0:30; 72°C for 0:30, the relative expression of target genes was calculated by 2^−ΔΔCT^. GAPDH was used as internal references in the quantitative analysis. The DNA products were also gel electrophoresis to determine their reliability. All primers were synthesized by Shanghai Sangon Bioengineering Co., LTD. The primers of each target genes as follow: mouse *KCNC2* Forward Primer: 5’-TCAATGGCACAAGTGTTGTTCT-3’, Reverse Primer: 5’-TAGCAGCTTTGGATGACAGCC-3’; mouse GAPDH Forward Primer: 5’-TGGCCTTCCGTGTTCCTAC-3’, Reverse Primer: 5’-GAGTTGCTGTTGAAGTCGCA-3’.

### Establishment of the drug-induced epileptic mouse model

To observe the effect of Kv3.2 on the susceptibility to epilepsy, models of epilepsy were established by pilocarpine 250mg/kg, PTZ 50mg/kg, penicillin 6mU/kg, KA 15mg/kg, respectively. Scopolamine (1mg/kg) was administered 30 min before pilocarpine intraperitoneal (IP) injection to reduce the undesirable peripheral cholinergic effects caused by pilocarpine. All the drugs were purchased from Aladdin (Shanghai Aladdin Biochemical Technology Co., Ltd., Shanghai, China). After intraperitoneal injection of the drug, all mice were continuously monitored and observed for 1 hour. The revised Racine Scale method was used to grade seizures(Racine, 1972).

### Preparation of acute brain slices, electrophysiology, single cell PCR detection in C57/6J mice

*Preparation of acute brain slices* The C57/6J mice were intraperitoneally injected with pentobarbital (50 mg/kg) and then decapitated. The brain tissue was quickly removed and placed in the dissection solution which contains (in mM) 92 NMDG, 2.5 KCl, 1.2 Na_2_HPO_4_, 30 NaHCO_3_, 20 glucose, 5 Na-ascorbate, 3 Na-pyruvate, 2 Thiourea, 10 MgCl_2_, 0.5 CaCl_2_ saturated with 95% O_2_ and 5% CO_2_ mixed gas in the state of ice-water mixture, coronal brain slices (200 μm) were prepared with vibratome (VT1200S, Leica). After sectioning, the slices were incubated in standard Artificial Cerebral Spinal Fluid (ACSF) which contains (in mM) 126 NaCl, 2.5 KCl, 1.25 Na_2_HPO_4_, 5 Na-ascorbate, 3 Na-pyruvate, 2 Thiourea, 2 CaCl_2_, 10 D-Glucose, and 26 NaHCO_3_ is oxygenated with gas mixture of 95% O_2_ and 5% CO_2_ for 30 min at 33°C. Then, the slices were allowed to recover for at least 30 min at room temperature before electrophysiology recording.

*Electrophysiology recording* For mice, whole-cell path-clamp recordings were performed using a multi-Clamp 700 B amplifier, with signals being recorded/analyzed using the Digidata 1550 B data acquisition system and pClamp 10.7 software package (Molecular Devices). The recording electrodes had a resistance of 3-6 mV when filled with the pipette solution containing (in mM) 100 K-gluconate, 50 KCl, 10 EGTA, 10 HEPES, 1 MgCl_2_ and 0.2 Na_2_-ATP. For recordings of native outward current in neurons, the extracellular solution was supplemented with tetrodotoxin (TTX, 0.001 mM) and CdCl_2_ (0.3 mM) to block sodium channels and calcium channel respectively. All in vitro recordings were made at 32-33°C. In the current clamp mode, the current threshold (rheobase) was evaluated with Ramp stimulation of 0-200 pA, and the firing frequency of action potential was evaluated at 200 pA which based on the resting potential level. Each scan last for 1 s and cells were clamped at normal RMP during recording.

*Single cell PCR detection* To determine the type of detected cells, after the electrophysiological test was completed, the detected cells were sucked into the recording electrode, and then the tip of the recording electrode was broken into the nested PCR reagent system (Prime ScriptTm RT reagent Kit with QDNA Eraser, Cat: RR0471, Takara), and a cDNA synthesis kit was used to synthesize cDNA following the manufacturer’s protocol (PrimeScriptm II1st Strand cDNA Synthesis Kit). The target genes were amplified by qPCR according to the instructions of kits (Cat.no. 638315 Takara, Japan), reversely. The DNA products were imaged using Quantum CX5 Gel Imager (Vilber Bio Imaging, France) after DNA agarose gel electrophoresis and used to determine the type of detected cells. GAPDH was used as internal references in the quantitative analysis. The PCR primers targeting genes are shown in Table 4. Reaction condition for nest PCR: 95°C for 2:00, 35 cycles of 95°C for 0:30, 55°C for 0:50, 72°C for 0:50; 72°C for 0:30. Reverse transcription PCR: 95°C for 2:00, 35 cycles of 95°C for 0:10, 58°C for 0:30, 72°C for 0:30; 72°C for 0:30.

### Cortical electroencephalogram (EEG) and intracranial EEG recording of hippocampal CA1 region in C57/6J mice

*Cortical EEG* Mice EEG recordings were performed after 3 weeks of the virus microinjection. Briefly, mice were anesthetized with pentobarbital (50 mg/kg), and implanted with recording electrodes under stereotaxic guidance. Cortical electrodes (Kedou Brain Computer Technology Co., Ltd, Suzhou, China) were implanted on the surface of the left temporal lobe and secured to the skull by super glue along with a ground electrode positioned over the epencephala. A week later, electrodes were connected to the EEG100C amplifier through bank cable and EEG recordings were performed. The signals were filtered (0.1 to 100 Hz) and digitized at 2 kHz. Time-frequency analysis and normalized power of EEG recordings [alpha (α), beta (β), gamma (γ)] were performed in the custom code and brainstorm based on MATLAB 2022.

*Intracranial EEG in hippocampal CA1 region* Briefly, intracranial electrode (Beijing Creation Technologies Co., Ltd, China) was implanted into the left dorsal hippocampus (AP: -2.50 mm; ML: +2.0 mm; DV: −2.0 mm) and secured to the skull by super glue along with a ground electrode positioned over the epencephala. Animals were allowed a week to recover. Electrodes were connected to the apollo amplifier through bank cable and used to recording cell discharges in hippocampal CA1 region. The signals were filtered (1 to 300 Hz) and digitized at 30 kHz. Time-frequency analysis and normalized power of EEG recordings were performed in the custom code and brainstorm based on MATLAB 2022.

### Statistics

MATLAB 2022 software was used to analyze EEG data, origin 2022 and graphpad 9 were used to analyze whole-cell patch recording data, and GraphPad prism 9 software was used for other data statistics. The unpaired t-test was used when data follows a normal distribution and has equal variances. If data follows a normal distribution but has unequal variances, the Welch’s t-test was used; if data does not follow a normal distribution, the Mann-Whitney U test was used. For comparing differences in proportions, the Chi-squared test was used.

## Data availability

All data are available in the main text or the supplementary materials. The mass spectrometry proteomics data have been deposited to the ProteomeXchange Consortium (http://proteomecentral.proteomexchange.org) via the iProX partner repository with the dataset identifier <u>PXD045397</u>.

## Acknowledgments

This study was supported by the following fundings: National Natural Science Foundation of China (81871075, 82071533) grants to HZ; National Natural Science Foundation of China (81870872) grants to XD; Science Fund for Creative Research Groups of Natural Science Foundation of Hebei Province (H2020206474); Basic Research Fund for Provincial Universities (JCYJ2021010); S&T Program of Hebei (21377732D) grants to JZ; Hebei Science and Technology Department - Hebei Medical University “Fourteen Five” clinical medical innovation research team support project (2022LCTD-A6) grants to WL. Natural Science Foundation of Hebei Province (C2019206583) grants to NZ.

## Author Contributions

Designing research studies: HLZ, WLL, NJZ.

Conducting experiments: YD, DZ, WG, YY, ATZ.

Acquiring data: YD, DZ, WG, NJZ, JD.

Analyzing data: NJZ, YD, DZ, WG, YY, ATZ, HRZ, JD.

Writing – original draft: NJZ, YD.

Writing – review & editing: HLZ, WLL, XND.

NJZ, YD, DZ contributed equally to this work.

